# Effect of the functional environment on the cardiac differentiation of iPSC

**DOI:** 10.64898/2026.01.08.695479

**Authors:** Nerea Pascual, Lara Rodríguez, Martxel Dehesa, Maurizio Prato, Nuria Alegret

## Abstract

Pristine carbon nanotubes (CNTs) have proven to be excellent supports for cardiac cell growth, survival and maturation, as well to improve cellular function, enhance spontaneous beating activity and benefit their cellular structure. Due to the large quantity of cardiomyocytes that have to be replaced for myocardial regeneration, iPSCs are the most promising candidates for robust generation of cardiomyocytes *in vivo/vitro*. Herein, iPSCs are cultured and differentiated into cardiomyocytes on functionalized carbon nanotubes (fCNTs). For this purpose, a first optimization of the type of plate and the number of iPSCs suitable for the passaging is performed. Thus, 5·10^5^ cells per cm^2^ are cultured in 24-well and 8-well plates. After 19-days of differentiation and maturation, calcium imaging was done to analyze the spontaneous beating behavior by means of beat frequency and amplitude, immunofluorescence was done to observe evaluate the degree of maturation by staining the sarcomere and the cell nucleus. A set of diverse functionalized CNTs were also tested: pristine CNT, ox-CNT, CNT-COOH, CNT-NH_2_, CNT-NO_2_ and CNT-SO_3_. Calcium analysis showed that all but the nitro-functionalization were beating, with acid-and oxygen derivative CNTs producing an increase in frequency with respect to control, while amino-functional groups decrease it. This suggests that the beating and contractile behavior of cardiomyocyte can be modulated according to the cardiac issue to be faced. In addition, CNT-SO_3_ produces a striated and elongated sarcomere, proper of the real tissue.

## 1. Introduction

The World Health Organization [1] reports that for the past 20 years, heart illnesses have been the top cause of death globally. More than 2 million individuals died between 2000 and 2019, reaching a peak of 9 million deaths in that year. In Europe, cardiovascular diseases, which include heart disease and stroke, account for more than 4 million annual fatalities, or 45% of all fatalities, and about 49 million individuals live with these conditions. Up until 2016, coronary heart disease and cerebrovascular disease were the leading causes of mortality from cardiovascular disease, while heart failure (HF) remained the major cause of morbidity and mortality worldwide [2]. On a chronic basis, the heart loses healthy cardiomyocytes (CM), leading to matrix degradation and fibrosis and, ultimately, HF. There is no cure for these frequent fatal disorders and current treatments include lifestyle changes and medicines to help relieve the symptoms. Because of that, and even under treatment, half of the patients with HF die after 5 years of diagnosis. The only definitive solution for the patients that are in the last stage of HF is cardiac transplantation, but availability of donor heart is a major limitation [3].

Even if prevention and treatments are available, there is a need to find a cure to HF, which would not only palliate the symptoms but also repair the damaged area. Considering the current difficulties in heart regeneration, along with the lack of promising remedial treatments and the urgent need to establish an advanced and effective therapy, tissue engineering and cardiac regeneration could be the remedy to these diseases. In that line, taking advantage of the system’s natural biology will allow for greater success in creating innovative and effective therapeutic treatment aimed at the replacement, maintenance, and repair of tissue and organ function [4].

Native cardiac muscle is characterized by its unique electrophysiological behavior, being capable of transferring electrical signals through CM couples with other heart cells, such as fibroblasts, smooth muscle cells, endothelial cells and macrophages. Myocardial regeneration or repair can only be achieved by the replacement of the approximately one billion CMs that die, for instance, during a myocardial infarction. Because of that, induced-pluripotent stem cells (iPSCs) are the most promising candidates for robust generation of large numbers of human cardiomyocytes *in vitro*, required for functional restoration of the failing heart. Ideally, cardiomyocytes differentiated from iPSCs with typical phenotype and genotype characteristics of a mature cardiomyocytes could be implanted directly into the damaged area. However, this has not yet been achieved in reality, and mainly due to the lack of cell retention after engraftment [5,6]. In addition, most of the reported iPSC-derived cardiomyocytes have an immature phenotype, resulting in low contractility and electrical instability [7,8]. For this reason, recent research suggests that engineering 3D scaffolds with similar properties to the native myocardial extracellular matrix (ECM) could offer a promising approach to support the survival of the transferred cells and to improve their maturation [9]. Castilho and co-workers reported a polycaprolactone cardiac patch created by melt electrowriting that yielded the best results of iPSC-derived cardiomyocyte cultures, as sarcomere density, alignment and length are improved. Furthermore, connexin 43 was present in all cardiac patches supporting electrophysiological coupling of iPSC-CMs [2]. Similarly, Jeong et al. also proposed a 3D-printed polycaprolactone based scaffold coupled with gold particle coated carbon nanofibers for patient specific cardiovascular therapies [10]. English et al. designed a micropatterned fibrin scaffold mimicking the native tissue structure. This micropattern was shown to promote cellular alignment in iPSC-CM cultures, with cells exhibiting enhanced cellular elongation, sarcomere alignment, connexin-43 staining and contractile properties [11].

Scaffolding substrates are essential not only for supporting the cells but also for their maturation. These scaffolds are artificial constructs that mimic the extracellular matrices and serve as temporary supports where individual cells are introduced to form new tissue [12]. As the ECM, the scaffold’s role is not only to define tissue boundaries, but also to support cell migration and contraction, delivery vehicle, matrix for cell adhesion, to facilitate certain cell process, provide a structural reinforce, prevent distortion and to be a barrier to infiltration [9]. In order to achieve the different characteristics of interest and to modulate the properties of the scaffolds, it is crucial to investigate between different types of biomaterials and shapes, thus providing the necessary electrophysiological properties. Among all, the suitable scaffold has to be biocompatible with the host tissue without inducing inflammatory or immune reactions [3].

Multiple experimental studies suggested that the use of nanoparticles could revolutionize the therapeutic frontiers of a huge number of disorders, such as neoplastic, heart or neurodegenerative diseases. Among conductive nanomaterials, carbon nanotubes (CNT) have numerous beneficial properties such as its unique electrical, mechanical and thermal properties [3]. Furthermore, their high mechanical strength and low weight is an added value to their conductivity and stability, making them a very interesting material for multiple fields, from delivery of pharmaceuticals and biosensors to biomedical applications [13]. More interestingly, CNTs can be functionalized so their chemical, physical and biological properties can be tunned and adapted to a specific application [3].

In addition, CNTs have also proven to have great potential for the treatment of cardiovascular diseases. More specifically, they have been used for diagnosis of atherosclerosis and myocardial infarction, to improve imaging modalities and characterization of atherosclerosis *in vivo* [3] and as a promising additive in an injectable hydrogel for cardiac tissue engineering applications [14]. CNT-based biomaterials have also been found to increase cardiomyocyte function and increase cardiomyocyte maturation [15,16]. The authors concluded in different *in vitro* studies that CNTs promote the formation of gap junctions, improve cardiomyocyte activity, and activate integrin pathways [13]. Furthermore, several studies demonstrate that local electrical stimulation using CNT as substrates favored and accelerated the healing processes of electroactive tissues, enhancing cell-cell and cell-substrate interactions [9]. Moreover, this fact was confirmed when interfacing CNTs with neuronal cells [17], which share several characteristics with cardiac cells, such as Ca2+ channels and electrical properties [18].

Herein, the hypothesis around which the present work was developed was that pristine and functionalized CNTs materials enhance the proliferation, growth and differentiation of cardiomyocytes derived from iPSCs. The presence of CNT will mean the addition of conductive features, *i.e.*, the key feature for electric stimulation and providing a new kind of interaction with biological systems. Hence, we evaluate diverse CNT-containing substrates in 2D and their functional effect on the growth, differentiation, properties, viability and beating behavior of iPSC-CM *in vitro* with the final aim to find an efficient substrate and the best conditions to promote cardiac regeneration.

## 2. Methodology

### Materials

Multi-walled carbon nanotubes (CNT) were purchased from Nanoamor Inc (Stock# 1237YJS: inner diameter 5−10 nm; outside diameter 20−30 nm; length 0.5−2 μm). Isoamyl nitrite and anilines (sulfanilic acid, 4-nitroaniline, 4-[(N-Boc)aminomethyl]aniline)) were purchased from Sigma Aldrich. The solvents were acquired from Carlo Erba Reagents SAS, Sabadell, Spain. All reagents and solvents were used as received with no further purification. All reagents and solvents were used as received with no further purification.

### Synthesis of functionalized CNT (f-CNT)

The functionalized CNTs, collected in **Error!**

**Reference source not found.**, were prepared as previously reported [19]. Briefly, was performed as follows.

#### Oxidation of CNT (ox-CNT)

p-CNT were dispersed in a concentration of 1mg/ml in a 3:1 solution of concentrated H_2_SO_3_ and concentrated HNO_3_. The solution was then sonicated for 6h, stirred overnight and diluted 1:10 in milliQ water. The ox-CNT were filtered over a PTFE membrane and rinsed with milliQ water until neutral pH was reached. Finally, the material was rinsed with ether and left to dry under vacuum overnight.

#### Tour reaction to synthesize CNT-COOH, CNT-SO_3_ and CNT-NH_2_

The tour reaction procedures to synthesize were adapted from reported literature [20]. Briefly, p-CNTs were dispersed in anhydrous DMF in a concentration of 1mg/ml and dispersed with the help of a sonication bath while purging the solution with Ar for 15min. 1.2 equivalents of the desired aniline were incorporated to the solution assuming the CNT samples to be of pure carbon. After equilibration of the system at 80°C, isoamyl nitrite (3.2 equiv per equivalent of aniline) was added dropwise to the reaction mixture and left under stirring at 80°C for 6h. The suspension was filtered, and the solid redispersed and washed with DMF until no color was observed in the filtered waters. The washing process was repeated as many times as necessary until a persistently colorless filtrate was obtained and continued in an identical manner with alternating cycles of Milli-Q water, and ethanol. Finally, the solid sample was rinsed with diethyl ether and left to dry at room temperature under vacuum. Specifically for CNT-NH_2_, cleavage of the Boc protecting groups was carried out suspending 4 in aqueous HCl solution (4M. 1mg/mL) and left to stir overnight at room temperature. The product was filtered over a PTFE membrane and rinsed with Milli-Q water until neutral pH value, rinsed with diethyl ether and dried under vacuum at room temperature.

### Preparation of the fCNT substrates

Bidimensional films of f-CNT were prepared by airbrushing aqueous dispersions of each f-CNT (1mg·ml^-1^) on glass coverslips (12mm #1, Thermo Scientific), in glass 8-well chamber (IBIDI) or P96 cell culture plastic plates, following 5 min annealing at 80 °C to ensure adherence. The spraying was performed from 8 to 10 longitudinal passes over the glass coverslip waiting for dryness of each layer in between passes. To speed up the drying process, the surface temperature of the substrate was kept around 50°C for the whole spraying procedure.

### Induced Pluripotent Stem Cells (iPSC) Culture

Human Induced Pluripotent Stem Cells (iPSC, cell line UCSD107i-2-6) were purchased from WiCell Research Institute, USA. The tissue origin of the iPSCs was skin fibroblast of a Caucasian European donor, a 74-year-old female without reported diseases. Culture, dissociation, thawing and freezing have been done following WiCell established protocols. Corning Matrigel® Phenol Red-Free (LOT: 4279010, 9,7 mg·ml-1, Invitrogen) was used as a coating matrix prior to the cell seeding to ensure cell attachment, mixed with IMDM (Iscove’s Modified Dulbecco’s Medium, Gibco). mTeSR™ Plus (mTeSR, Stem Cell) was used for iPSC medium and mixed with ROCK inhibitor (Stem cell, Y-27632 RHO/ROCK pathway inhibitor; Inhibitor ROCK1 and ROCK2) when passaging or thawing. All cell passages were done when the colonies reach a confluency between 60-80% using ReLeSR (Stem Cell) or Accutase (Stem Cell). Accutase was only used when cell counting was needed since it lifts and separates the cells into single cells, by digesting integrins and cadherins. ReLeSR, instead, detaches the cells without breaking the aggregates, which avoid single cells and debris formation. For freezing, 500μl of Cryostor (Stem Cell) per tube of cryovial was used.

### iPSC differentiation into cardiomyocytes

The STEMdiff Cardiomyocyte Differentation and Maintenance Kit (Stem Cell) was used to generate cardiomyocytes (cardiac troponin T-positive [cTnT+]), derived from a clump culture of iPSCs. The protocol lasted from 15 to 18 days: it took 8 days to differentiate the iPSC into cardiomyocytes (iPSC-CM), and at least 7 days more to reach full maturity. It was expected the cardiomyocytes to beat between day 12 and day 18. Afterwards, the resulting beating iPSC-CM were used in various downstream applications and analyses.

### Viability assay

The viability of cells grown on the scaffolds was evaluated with the modified LDH CytoTOX96 Nonradioactive Cytotoxicity Assay kit (Promega). The CytoTox 96 Assay quantitatively measured lactate dehydrogenase. The viability protocol is briefly summarized as follows: 30μl of lysis Solution (10X) [9% (v/v) Triton® X-100 in water] were added to each well, 15μl per 100μl of culture medium, the plate was incubated for 30 minutes at −80°C and 20 minutes at 37°C. Then the culture of each well was transferred into an Eppendorf and centrifuged at 10^3^ RCF for 10 minutes at 4°C to separate de CNT. 50μl of supernatant were transferred to each well of a fresh 96-well flat-bottom (enzymatic assay) plate. 50μl of CytoTox Reagent were added to each well of the enzymatic assay plate. The plate was protected from light and incubated for 30 minutes at room temperature. Finally, visible wavelength absorbance data was recorded at 492nm within 1 hour after adding 50μl of Stop Solution using a standard 96-well plate reader in the multi-mode microplate reader (BMG Labtech), for luminescence, fluorescence and fluorescence anisotropy assays.

### Calcium Imaging

Intracellular calcium signaling of iPSC-CM after 14-18 days of culture was recorded to assess the electrical activity of CMs growing on the 2D materials and controls. Cell-permeant fluo-4 AM was added to each sample and incubated for 15 min, according to manufacture instructions. Samples were then washed three times with warm media and, subsequently, calcium transients imaging during spontaneous beating of iPSC-CMs were recorded for 90 s. All recordings were done with an Inverted ECLIPSE Ti-S / L100 fluorescence microscope (Nikos) using the FITC filter. Experiments were performed in duplicate and averaged. Approximately 10 videos per well were recorded. The amplitude and frequency of calcium signals were later analyzed. For the analysis of the results the mean and standard deviation were obtained. For statistical analysis, outliers were identified using ROUT method with a Q of 0,5%, normality and lognormality tests were done and based on those results, rather an unpaired t-test or a Mann-Whitney test was performed using the GraphPad Prism 8 (GraphPad Software, Inc., San Diego, CA., EEUU). To take into account the signals, two requirements were followed: (i) the peaks shall be constant in both peak width and amplitude; (ii) the signals should have typical morphology, thus fast rise and decay kinetics with no oscillations. Moreover, the discard criteria were used with single-peak signals, signals with peaks with amplitudes less than 15% of the maximum amplitude and signals with irregular phase.

### Immunocytochemistry

Immunocytochemistry was carried out after 19 days of culture to assess cardiomyocyte maturation and culture purity. To this end, cells were stained for the cardiac-specific marker α-sarcomeric actinin (primary antibody: 1:200, Sigma; secondary: Alexa Fluor 647-conjugated goat anti-rabbit, 1:200, Invitrogen). Gap junction distribution between the cardiomyocytes was evaluated concurrently using an anti-Connexin 43 antibody (1:200, Sigma), with a subsequent Alexa Fluor 488-conjugated goat anti-mouse secondary antibody (1:200, Invitrogen).

The staining protocol proceeded as follows: Cells cultured on coverslips were initially washed with PBS and then fixed using 4% paraformaldehyde (PFA) in PBS for 15 minutes. Permeabilization with 1% Triton X-100 followed for 1.5 hours, and non-specific binding was blocked with 2% BSA in PBS for 1 hour. Primary antibodies were applied and incubated overnight. The cells were then incubated with the secondary antibodies for 1 hour, and the nuclei were counterstained with DAPI for 2 minutes. Fluorescent images were captured from five distinct regions per sample (n=4) using LS90 Zeiss confocal microscopy at 10X, 20X, and 40X magnifications, maintaining consistent instrument settings throughout the experiment.

### Images evaluation and statistical analysis

For immune staining analysis, data was collected in quadruplicates from two different experiments. For calcium signals, data was collected and analyzed from at least 10 fields per sample and averaged.

Calcium measurements are reported as the fractional change in fluorescence intensity relative to baseline (F/F0), which was determined as follows. Within a temporal sequence of fluorescence images, a region of interest (ROI) was drawn around a portion of each cardiomyocyte to be analyzed. The fluorescence signal from each terminal was calculated as the pixel-averaged intensity within each ROI, yielding the common sinusoidal beating plot represented. The absolute minimum value of the pixel averaged intensity from each of these ROIs in the whole temporary region was taken as the baseline fluorescence (F0) for that cell. At each time point, the fluorescence value was rationed against the baseline value to yield F/F0. The amplitudes of Ca^2+^ transients were determined by the maximum values of the beating peaks; the frequency of the Ca^2+^ transients were calculated by the time differences between adjacent peaks. The precise boundary of each ROI was kept constant between all the analyzed beating cells.

Data distribution was evaluated with Shapiro-Wilk normality test (GraphPad Prism 8). Statistical significance between experimental groups for calcium imaging was determined using student t-test analysis or one-way ANOVA followed by Tukey’s post-hoc test for multiple pairwise comparisons. In all cases, adjusted *P* values of less than 0.05 were considered statistically significant.

## 3. Results and Discussion

Even though the commercially available differentiation kit for iPSC-CM is designed to yield highly mature and purely differentiated cardiac myocytes, optimal results necessitated the optimization of specific parameters. Therefore, we have studied the seeding density, the functional group attached to the CNT-substrates and the conductivity of the substrates. We specifically aimed to achieve a cardiomyogenic phenotype, ensuring complete differentiation into a mature cardiomyocyte lineage. This phenotype was evidenced by the culture’s uniform structure, beating behavior, contractility, and the presence of striated and elongated sarcomeres.

### 3.1 Effect of the seeding density in the cardiac differentiation

One of the main parameters for a successful growth and differentiation of iPSC into cardiomyocytes is the initial seeding density. We first studied this effect on matrigel®-coated (MG) and matrigel®-CNT-coated (MG-CNT) 24-well plates, and compared cultures with initial seeding of 5·10^5^ and 8.5·10^5^ cells per well.

A first characteristic of a mature functional cardiomyogenic tissue is the uniform beating. To assess the functionality and maturation of the iPSC-CM, we have first analyzed spontaneous intracellular calcium transients after 19 days of differentiation. As shown in Figure 2, amplitude demonstrated a positive correlation with increasing seeded cell numbers on both MG and MG-CNT substrates. A notable observation was the generally lower amplitude in the presence of CNT for lower seeding densities. For instance, with 5·10^5^ seeded cells, the amplitude values for MG (1.1 ± 0.1) and MG-CNT (1.0 ± 0.5) were statistically different. Following the same trend, at 8.5·10^5^ seeded cells, a significant increase in amplitude was observed, rising from 1.2± 0.2 on MG-coated substrates to 1.5± 0.4 on MG-CNT-coated substrates. Conversely, the period was significantly higher at the lowest seeding density across all substrates, with similar values observed for both MG and MG-CNT coatings at higher seeding densities. Specifically, for 5·10^5^ seeded cells, the periods were 4.2 ± 2.1s on MG and 3 ± 1.2s on MG-CNT. In contrast, for 8.5·10^5^ seeded cells, the periods were 2.1 ± 0.2s and 2.5 ± 1.4s, respectively. This inverse relationship between seeding density and period indicates that lower cell numbers result in fewer beats per minute (BPM), consequently leading to reduced calcium flux within the cardiomyocytes. These findings confirm that a greater number of seeded cells enhances contraction behavior, while the contribution of CNTs appears to primarily influence amplitude, not beat period.

**Figure 1.**
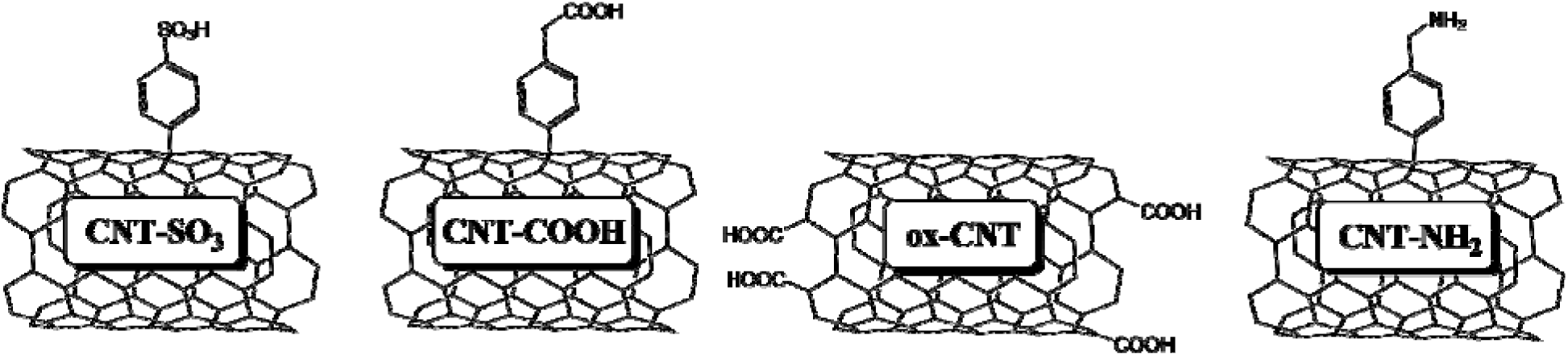
Scheme of functionalized CNTs, CNT-SO3, CNT-COOH, ox-CNT and CNT-NH2.

**Figure 2.**
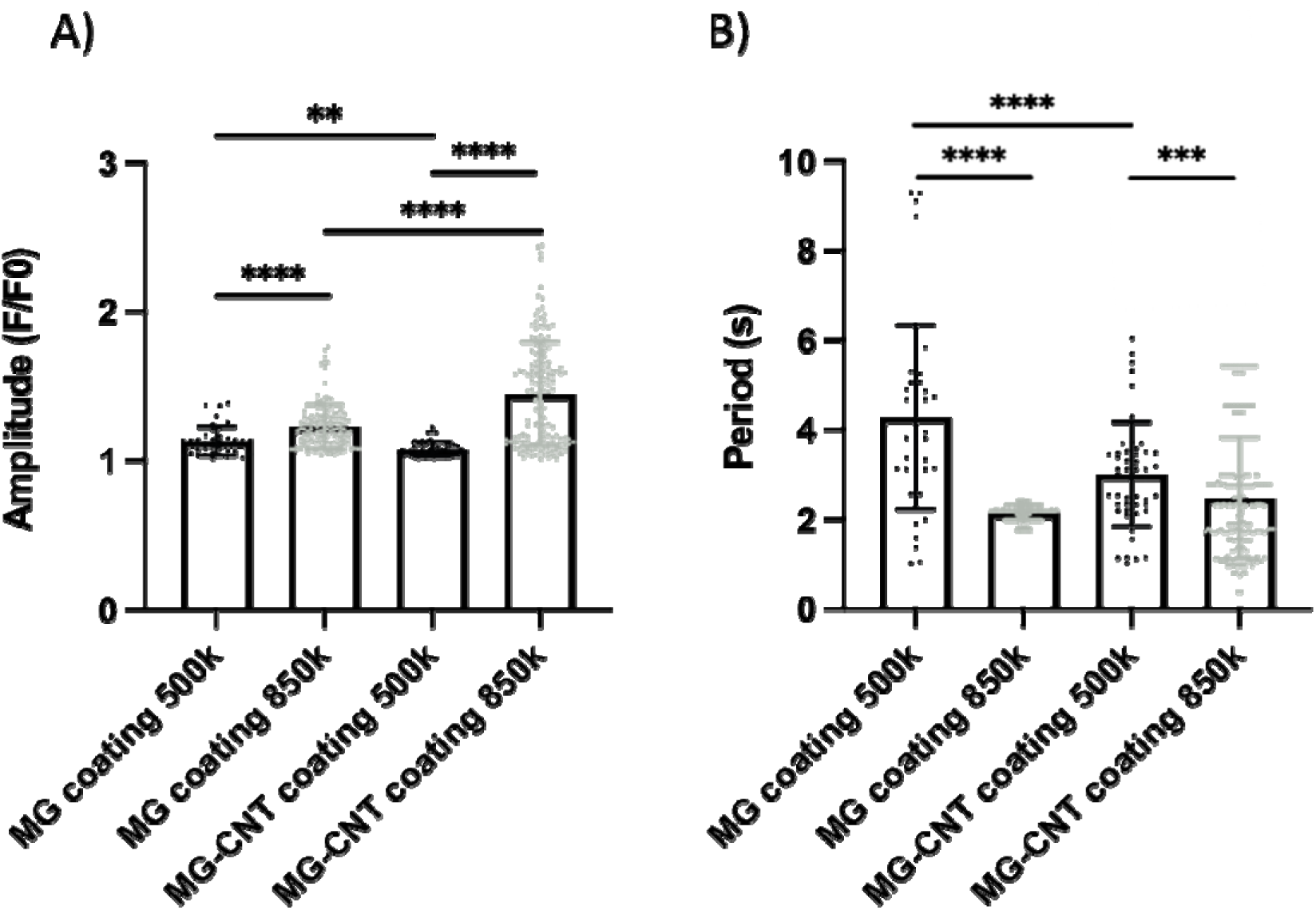
**A)** Comparison of amplitude and **B)** period of calcium intramolecular transient signals of cells seeded in a P24 well plate at 5·10^5^ cells and 8.5·10^5^ cells density per well with and without CNT. P-value not significant, >0.005, in the case of amplitude and significant, <0.005, in the case of period (Mann-Whitney test).

The striated and elongated sarcomere as well as the number and organization of gap junctions between differentiated cells are also two features of mature cardiac tissue. Thus, immunofluorescence was performed to assess the organization and phenotype of the iPSC-CM. Two specific markers were employed: sarcomeric α-actinin to stain the sarcomere, specific for cardiomyocytes, and in 8.5·10^5^ cells seedings also connexin-43 (Cx-43) was used to stain the cell-cell gap junctions. The images collected in Figure 3 show a clear difference in the sarcomere structure and organization between seeding density, but more significantly between MG and CNT coatings. At first glance, in either the coatings, the iPSC-CM seeded in large density - 8.5·10^5^ cells - show a tissue-like structure after differentiation, while in low cell-content seeding individual cells are found, being rounder and smaller. Furthermore, in 5·10^5^ cell seedings, the sarcomere is located mainly on the cell walls, what make cells clearly distinguishable, and no striation observed. Interestingly, only iPSC-CM differentiated on CNT coatings in the large seeding density, thus 8.5·10^5^ cells, show a sarcomere that is elongated inside the cell, although it is not homogeneous nor striated. Finally, Cx-43 staining reveals that the presence of CNT yields to a more interconnected structure. Overall, these results demonstrate that the number of cells and the coating influences the subsequent growth, expression and maturation of differentiated cardiomyocytes.

**Figure 3.**
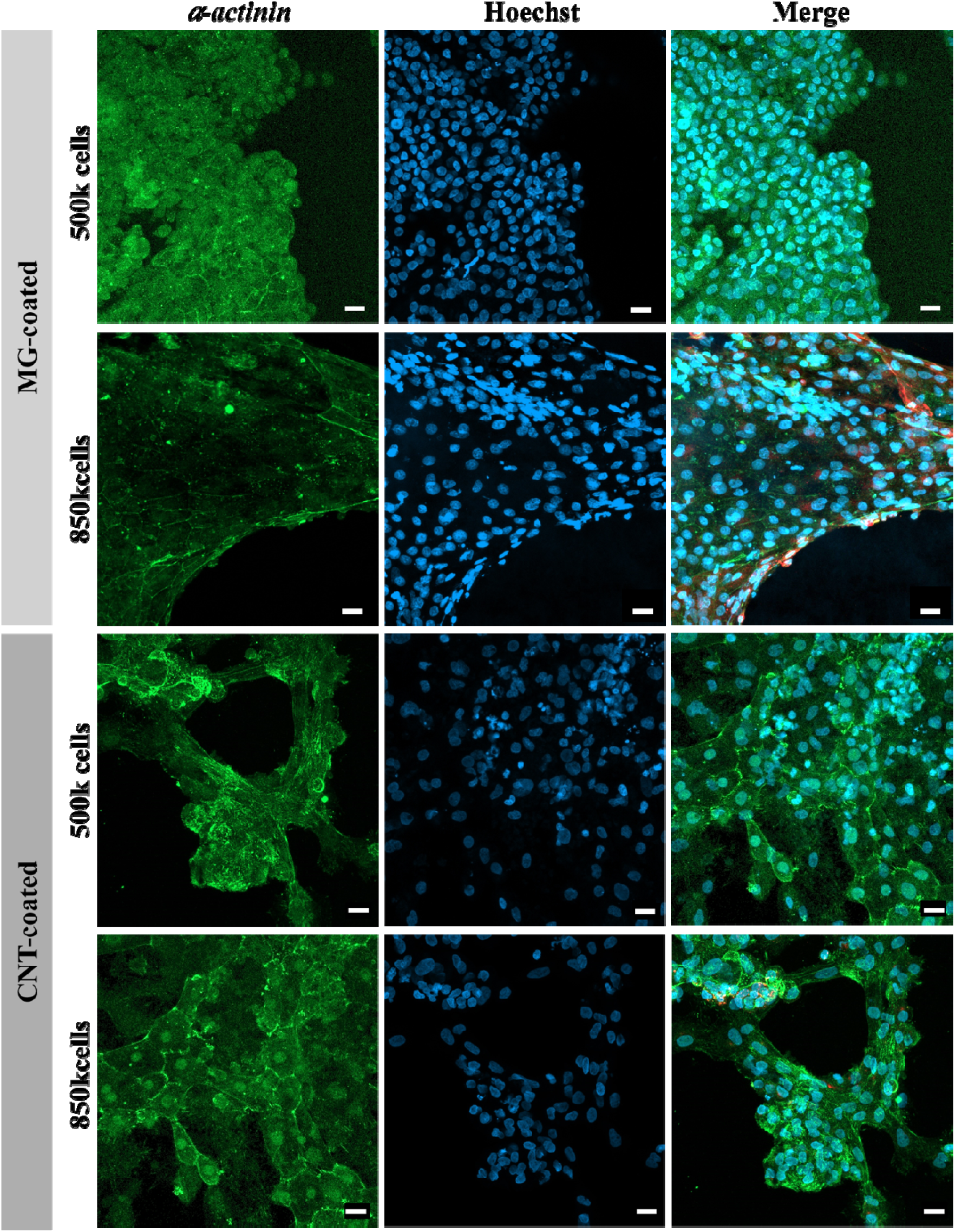
Sarcomeric α-actinin (green), Hoechst nuclei stainning (blue) and merge immunofluorescence images of cardiomyocytes seeded on MG- and CNT-coatings, with 5·10^5^ cells and 8.5·10^5^ cells seeded per well. Scale bars: 20µm.

### 3.2 Effect of the functional group in the differentiation

As demonstrated in the previous section, CNT improve the behavior, beating and phenotype of cardiomyocytes when seeding 8.5·10^5^ cells per P24-well, although the cells do not show an ideal mature uniformly striated sarcomere. For this reason, we aim to elucidate whether the incorporation of specific functional groups on the surface of the CNT might yield to a more mature phenotype of the differentiated iPSC-CM cells. Several functionalization were prepared to produce functionalized CNT (fCNT) with different functional groups and surface charge (see Figure 1): oxidized CNT (ox-CNT), enriched of negatively charged groups; the acid-terminated CNT (CNT-COOH); the positively charged amine-terminated CNT (CNT-NH_2_); and the negatively charged sulfonic-terminated CNT (CNT-SO_3_). All materials were airbrushed on 12 mm coverslips, UV-sterilized, and seeded with 8.5·10^5^ cells per well inside P24 well-plates. It is worthy to note that nitro-terminated CNT (CNT-NO_2_) materials were also synthesized, but since no beating of differentiated iPSC-CM was observed during the differentiation, this material was discarded and not further characterization was done in posterior analyses. Matrigel and pristine CNT coatings were used as controls.

Following differentiation, beating iPSC-CMs were observed in all materials. The strongest, most homogeneous, and most rhythmical beating was seen in the CNT and CNT-SO_3_ wells under the optical microscope. Calcium analyses, presented in Figures 4b-c and Table S1, show that all materials initially exhibit a homogeneous and rhythmic beating behavior. Interestingly, the negatively charged CNT- SO_3_ and the electron-attracting groups of ox-CNT and CNT-COOH show less intense and faster beating, while the positively charged CNT-NH_2_ substrate displays the highest amplitude and lowest beating, resulting in the largest period. As expected, the pristine CNT exhibits amplitude and period values that fall between those of the two charged groups, but with values closer to the electron-acceptor groups (oxidized, acid, and sulfonic groups). This is consistent with the well-known electron-acceptor nature of pristine CNTs [21]. Furthermore, statistical analysis reveals significant differences between the materials, except for CNT-SO_3_ and pCNT (Figure 4b). These findings underscore the importance of the materials’ conductive properties, and especially the type of functional group, in the beating behavior and functionality of differentiated iPSC-CMs. For instance, the similarity in the effect of CNT-COOH and ox-CNT on cardiomyocytes is consistent with both being uncharged materials with a high oxygen content and a negative electron density displaced toward their functional groups. Conversely, the CNT-SO_3_ and CNT-NH_2_ substrates, which are oppositely charged at a physiological pH of 5 (negative and positive, respectively), show divergent effects: while CNT-SO_3_ has no effect on beating, CNT-NH_2_ decreases the beating frequency when compared to control.

**Figure 4.**
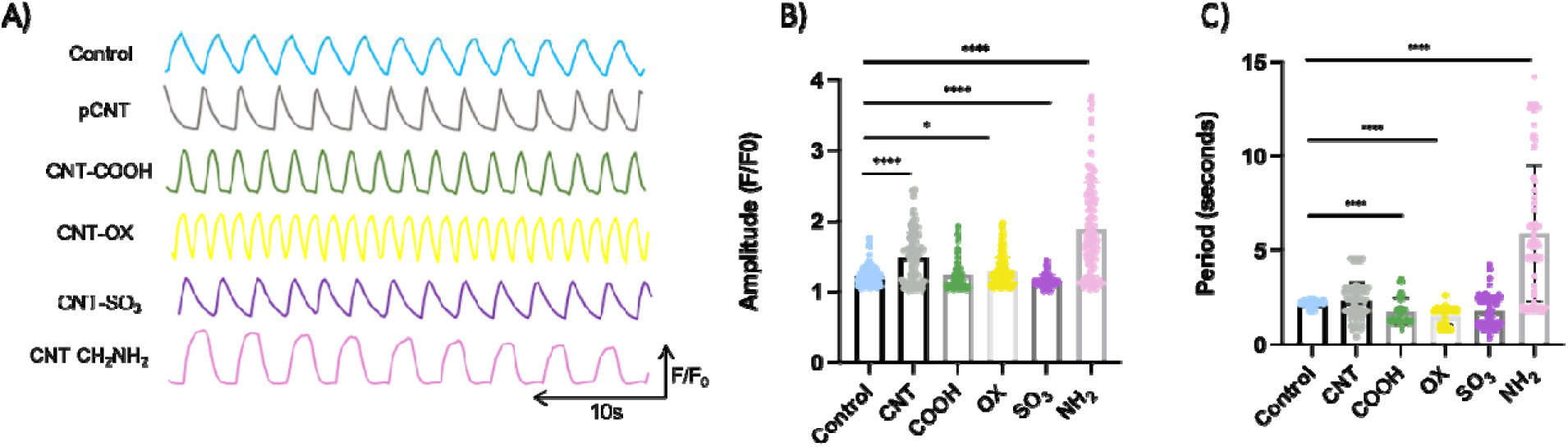
**A)** Calcium imaging plot of the cardiac cell spontaneous beating behavior of cultured iPSC derived CMs. B) Comparison of calcium signal amplitude and C) period between control and fCNT. P value significant, <0.0.05. None of the materials followed a normal distribution so Mann-Whitney tests were performed.

Finally, we performed immunofluorescence to evaluate the mature structure of the sarcomere and the tissue. Differentiated cells were stained with sarcomeric α-actinin and Cx43 after 15–18 days of differentiation. A representative set of images is displayed in Figure 5. Among all tested materials, and in agreement with the strong beating observed under the optical microscope, cell seeded on CNT-SO_3_ stood out. They exhibited the ideal mature cardiac tissue phenotype, characterized by homogeneously striated and elongated sarcomeres, similar to native tissue. In this case, single cells were not visible, and the nuclei were among the largest within all the materials (see Figure S1). The remaining materials showed a similar phenotype to the CNT substrates, forming cardiac tissue with sarcomeres primarily accumulated in the cell walls. Interestingly, the nuclei differed between materials, being the most round and small in CNT-COOH and more elongated in CNT-NH_2_ and ox-CNT, which suggests the formation of enlarged and elongated cells on these two substrates. Furthermore, Cx43 was only observed surrounding the cell walls in CNT-COOH, demonstrating connectivity between neighboring cardiomyocytes and, thus, their capacity for intercellular communication. In all other materials, Cx43 was primarily displayed within the cellular cytoplasm.

**Figure 5.**
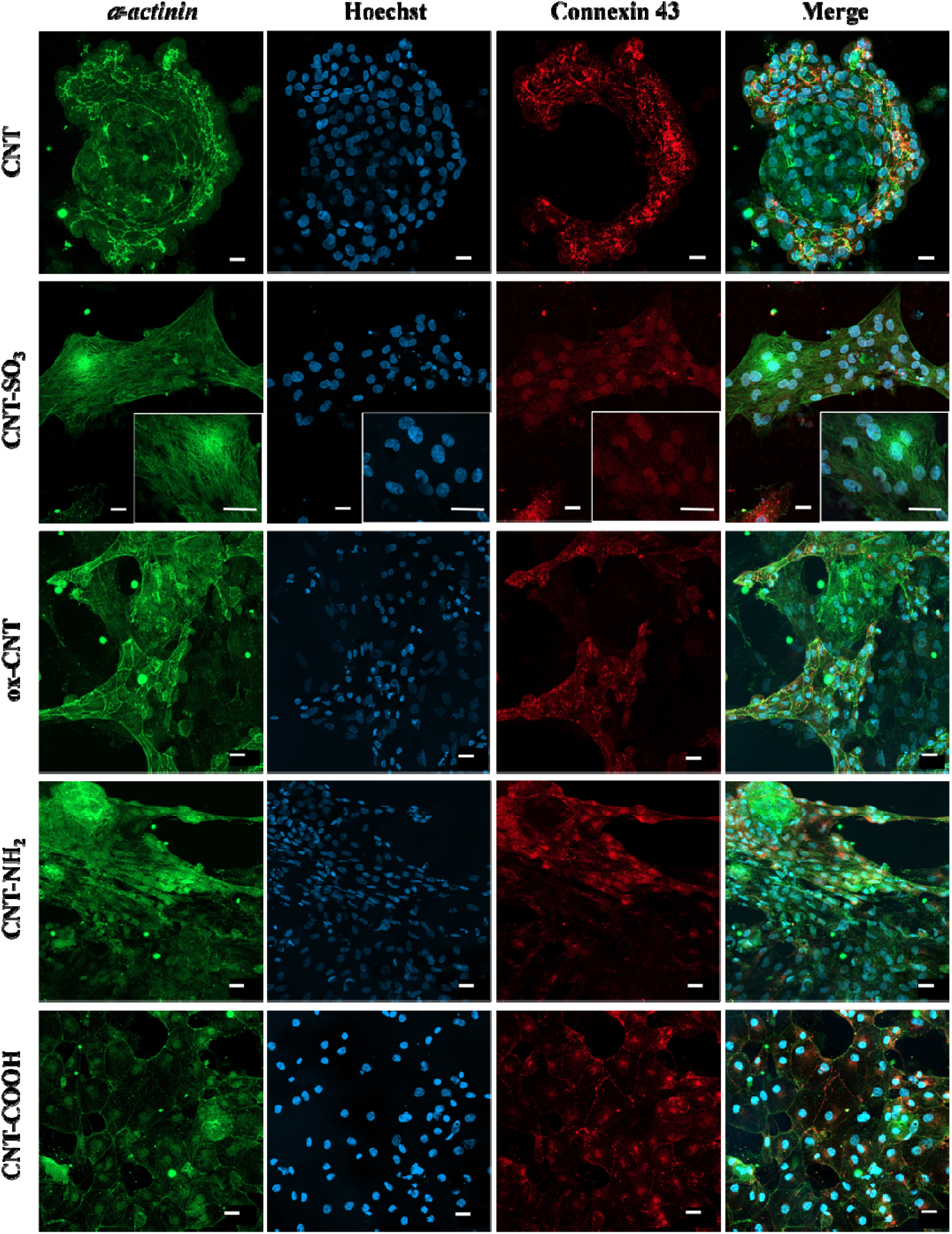
Sarcomeric α-actinin, Hoechst, connexin 43 and merge immunofluorescence images. Images of control, pCNT, CNT-SO_3_, ox-CNT, CH_2_NH_2_ and CNT-COOH, cardiomyocytes at 20X are shown. Images of CNT-SO_3_ are also shown at 40X. Scale bars: 20µm.

### 3.3 Effect of the conductive material in the iPSC-CM differentiation

To better understand the role of electrical conductivity in cardiomyocyte differentiation and culture, we also evaluated other widely used conductive polymers and their combinations with CNTs. Coatings were prepared by airbrushing aqueous dispersions of CNTs, polypyrrole, (PPy, smashed from a 3D scaffold prepared as reported [12], PPy/CNT, poly(3,4-ethylenedioxythiophene) polystyrene sulfonate (PEDOT/PSS), PPy/PSS and PEDOT/CNT on glass coverslips until a homogeneous layer was formed. The coverslips were then placed into 24-well plates, and 5 x 10^5^ iPSCs were seeded per well for differentiation. Matrigel coating served as control.

*In vitro* viability was analyzed using a modified lactate dehydrogenase (LDH) assay. According to the results plotted in Figure 6, no cytotoxicity was observed for any of the materials except for PPy/PSS. This lack of viability for PPy/PSS may be due to poor cell adhesion and/or proliferation, effectively ruling it out as a candidate material for cardiac regeneration. It is noteworthy that the PEDOT/CNT substrate showed exceptionally high viability, almost double the average of the other materials. The remaining materials also demonstrated higher viability than the control, suggesting that conductive materials, in general, enhance iPSC growth. However, because none of these materials, except for CNT, showed spontaneous beating during differentiation, calcium analyses were not performed.

**Figure 6.**
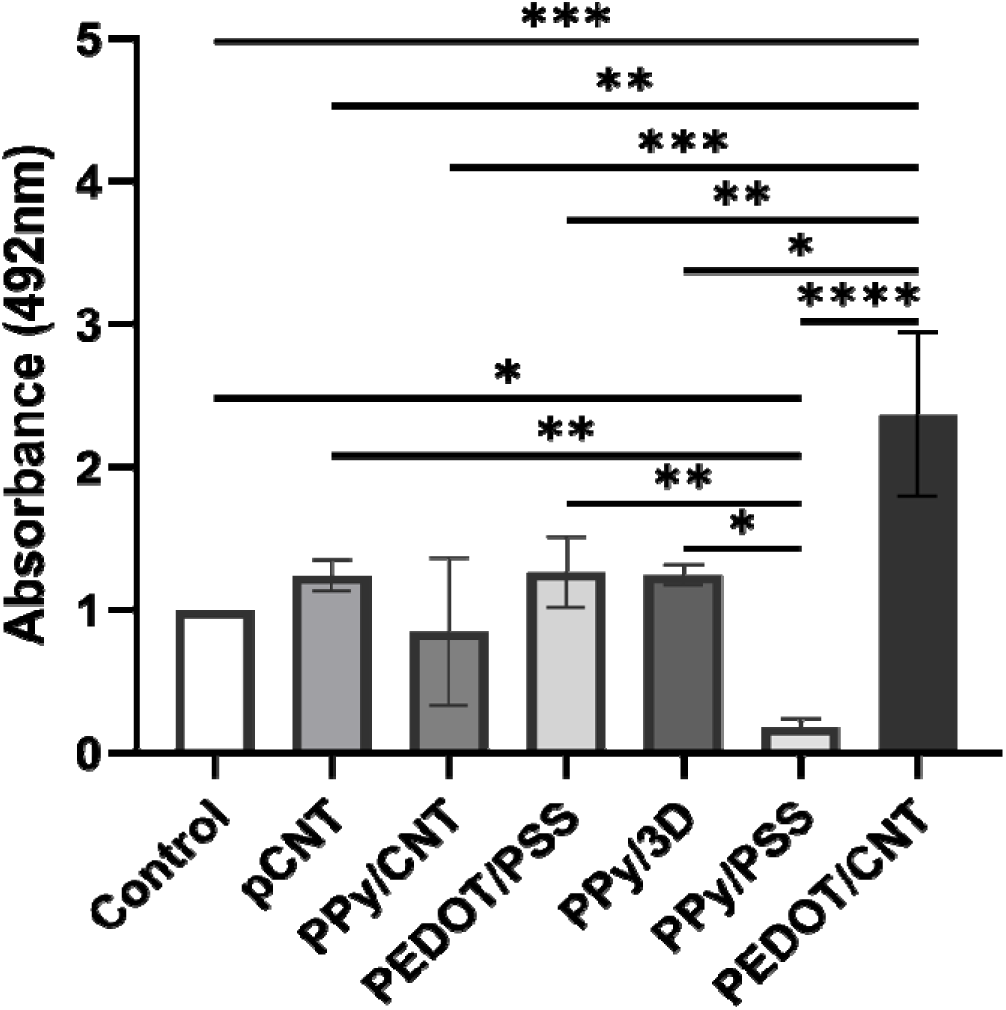
Viability assay of iPSC-derived cardiomyocytes cultured on different conductive materials 2D coatings mean ± SEM.

Despite the absence of spontaneous beating in the differentiated iPSC-CMs on the PPy- and PEDOT-based substrates, immunocytochemistry was used to assess their level of maturation. The images collected in Figure 7 show positive staining for the cardiac marker sarcomeric α-actinin in all PEDOT-based materials, as well as in PPy and PPy/CNT. However, the sarcomeres lacked a striated structure and were primarily localized along the cell membranes. This observation indicates some degree of maturation but suggests a non-ideal tissue formation.

**Figure 7.**
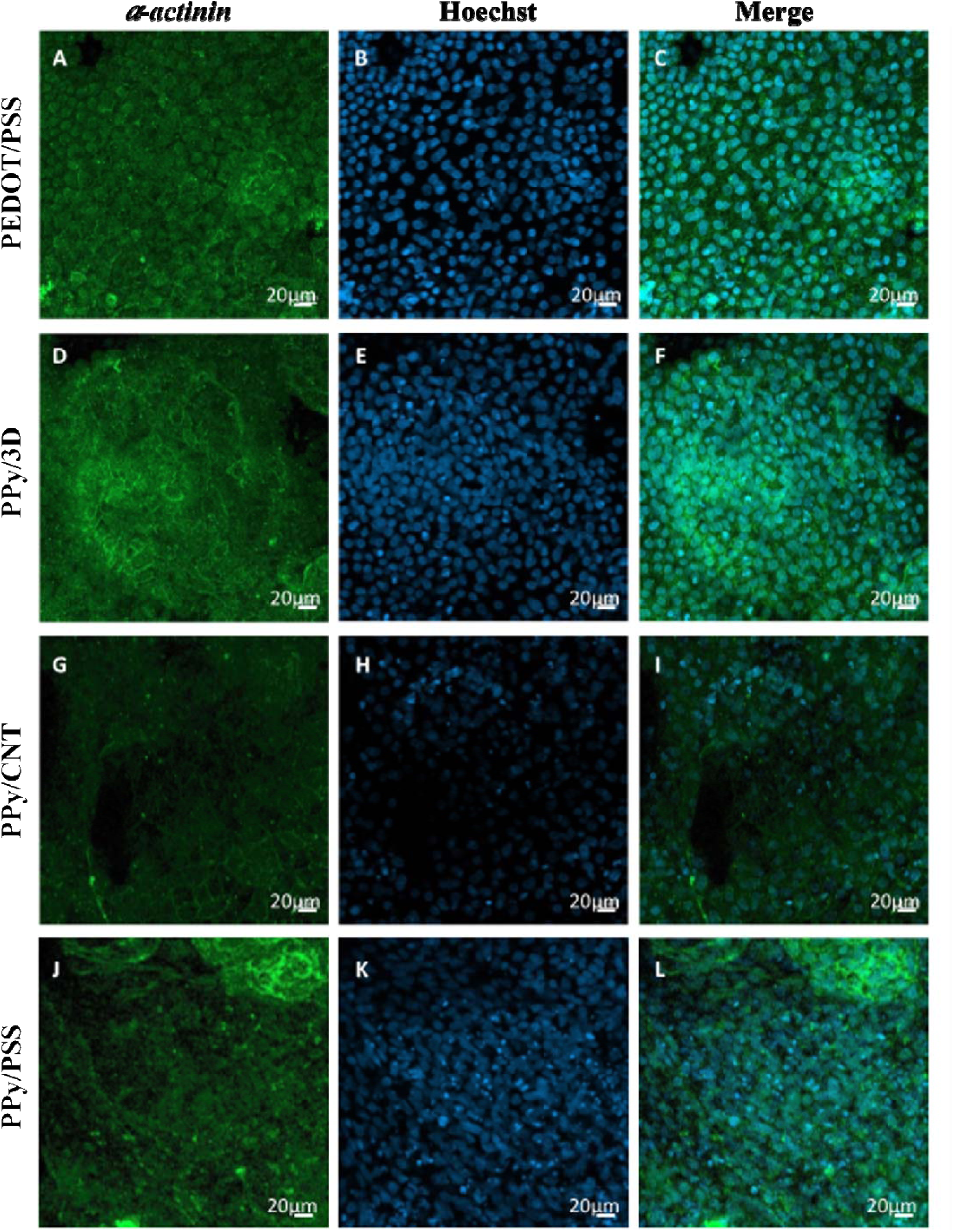
Sarcomeric α-actinin, Hoechst and merge immunofluorescence at 20X magnification. Panels A, B and C show control cardiomyocytes in presence of PEDOT/PSS. D, E and F panels show cardiomyocytes in the presence of PPy/3D. G, H, and I panels show cardiomyocytes in the presence of PPy/CNT. Panels J, K and L show cardiomyocytes in presence of PPy/PSS. Scale bars: 20µm.

## 4. Conclusions

Given the need for alternatives to current cardiovascular treatments, tissue engineering has emerged as a field with significant potential. Cardiomyocyte-derived iPSCs are promising candidates for regenerative therapy, although their immaturity presents a major challenge. Consequently, identifying a suitable material or scaffold that induces their maturation and promotes spontaneous beating is a crucial step toward solving this problem.

This work has demonstrated the importance of optimizing all culture variables for cardiomyocyte differentiation. It was shown that cardiomyocytes can grow and differentiate in P24 well-plates, and that a seeding density of 5 x 10^5^ cells per cm² is sufficient for proper proliferation and expression of the CM phenotype. Having established the potential of CNTs, our results demonstrate the distinct effects of nanotube functionalizations. It was observed that materials such as CNT-COOH, ox-CNT, and CNT-SO_3_ increased the number of beats per minute compared to the control, with CNT-SO_3_ standing out for its particularly notable effect. Conversely, pCNT and CNT-NH_2_ decreased the beating rate. These results underscore that not only conductivity but also the electrical charge and chemical properties of the functional groups directly influence cardiomyocyte behavior.

In addition, a strong correlation is observed between the quality of tissue structure (striated sarcomeres, elongated cells) and the functional performance of the cells (strong, rhythmic beating). The CNT-SO_3_ is the clearest example, as its ability to promote an ideal tissue structure is directly associated with the strongest beating. This reinforces the idea that structural maturation and electrical functionality are intrinsically linked. The diverse responses of the CMs to these functionalizations suggest a powerful tool for controlling specific cellular parameters, allowing for the selection of a material based on the targeted cardiac problem. For example, materials that increase the beating period could prove beneficial in the treatment of bradycardias. Finally, this pioneering work reveals the complexity of cellular maturation and underscores the need to validate our 2D material findings for their application in 3D scaffolds. The fact that PPy-and PEDOT-based materials did not exhibit spontaneous beating in 2D, despite cellular viability, suggests that full functionality requires specific scaffolding conditions beyond simple viability and marker expression. Therefore, future efforts should focus on establishing standardized sterilization protocols and creating robust methods for monitoring cardiomyocyte differentiation within the 3D matrix. Ultimately, this pioneering research serves as a vital starting point for developing 3D materials and, consequently, advanced medical devices like conductive and stretchable patches for cardiac diseases, which could revolutionize therapeutic approaches.

## Author Contributions

The manuscript was written through contributions from all authors.

## Data availability

The datasets generated during and/or analysed during the current study are available from the corresponding author on reasonable request.

## Consent to Publish declaration

Not applicable

## Consent to Participate declaration

Not applicable

## Ethics declaration

Not applicable

## References

1. World Health Organization. Cardiovascular-Diseases-(Cvds) @ Www.Who.Int. (2017).

2. Castilho, M. et al. Melt Electrowriting Allows Tailored Microstructural and Mechanical Design of Scaffolds to Advance Functional Human Myocardial Tissue Formation. Adv Funct Mater 28, (2018).

3. Peña, B. et al. Carbon Nanotubes for Cardiac Applications. in Carbon Nanostructures for Biomedical Applications 223–256 (The Royal Society of Chemistry, 2021). doi:10.1039/9781839161070-00223.

4. Nerem, R. M. & Schutte, S. C. The Challenge of Imitating Nature. in Principles of Tissue Engineering 9–24 (Elsevier, 2014). doi:10.1016/B978-0-12-398358-9.00002-1.

5. Jiang, X. et al. Maturation of pluripotent stem cell-derived cardiomyocytes: limitations and challenges from metabolic aspects. Stem Cell Res Ther 15, 354 (2024).

6. Sugiura, T., Shahannaz, D. C. & Ferrell, B. E. Current Status of Cardiac Regenerative Therapy Using Induced Pluripotent Stem Cells. Int J Mol Sci 25, 5772 (2024).

7. Strimaityte, D. et al. Contractility and Calcium Transient Maturation in the Human iPSC-Derived Cardiac Microfibers. ACS Appl Mater Interfaces 14, 35376–35388 (2022).

8. Sottas, V. et al. Improving electrical properties of iPSC-cardiomyocytes by enhancing Cx43 expression. J Mol Cell Cardiol 120, 31–41 (2018).

9. Wu, P. et al. Maturation strategies and limitations of induced pluripotent stem cell-derived cardiomyocytes. Biosci Rep 41, (2021).

10. Jeong, Y.-J. et al. 3D-printed cardiovascular polymer scaffold reinforced by functional nanofiber additives for tunable mechanical strength and controlled drug release. Chemical Engineering Journal 454, 140118 (2023).

11. English, E. J., Samolyk, B. L., Gaudette, G. R. & Pins, G. D. Micropatterned fibrin scaffolds increase cardiomyocyte alignment and contractility for the fabrication of engineered myocardial tissue. J Biomed Mater Res A 111, 1309–1321 (2023).

12. Alegret, N. et al. Three-Dimensional Conductive Scaffolds as Neural Prostheses Based on Carbon Nanotubes and Polypyrrole. ACS Appl Mater Interfaces 10, 43904–43914 (2018).

13. Ghai, P., Mayerhofer, T. & Jha, R. K. Exploring the effectiveness of incorporating carbon nanotubes into bioengineered scaffolds to improve cardiomyocyte function. Expert Rev Clin Pharmacol 13, 1347–1366 (2020).

14. Peña, B. et al. Biocompatibility Assessment of an Injectable Carbon Nanotube-Functionalized Reverse Thermal Gel for Cardiac Tissue Engineering Applications. ACS Appl Bio Mater 8, 4743–4755 (2025).

15. Martinelli, V. et al. Improving cardiac myocytes performance by carbon nanotubes platforms†. Front Physiol 4, (2013).

16. Barrejón, M., Marchesan, S., Alegret, N. & Prato, M. Carbon nanotubes for cardiac tissue regeneration: State of the art and perspectives. Carbon N Y 184, 641–650 (2021).

17. Pampaloni, N. P. et al. Sculpting neurotransmission during synaptic development by 2D nanostructured interfaces. Nanomedicine 14, 2521–2532 (2018).

18. Alano, C. C. et al. Differences among cell types in NAD ^+^ compartmentalization: A comparison of neurons, astrocytes, and cardiac myocytes. J Neurosci Res 85, 3378–3385 (2007).

19. Daou, B. et al. Organic Functional Group on Carbon Nanotube Modulates the Maturation of SH SY5Y Neuronal Models. Macromol Biosci 23, (2023).

20. González-Domínguez, J. M. et al. Effect of Various Aminated Single-Walled Carbon Nanotubes on the Epoxy Cross-Linking Reactions. The Journal of Physical Chemistry C 115, 7238–7248 (2011).

21. Vardakas, P. et al. Pristine, carboxylated, and hybrid multi-walled carbon nanotubes exert potent antioxidant activities in in vitro-cell free systems. Environ Res 220, 115156 (2023).

