## Supplementary Information for "Effect of the functional environment on the cardiac differentiation of iPSC"

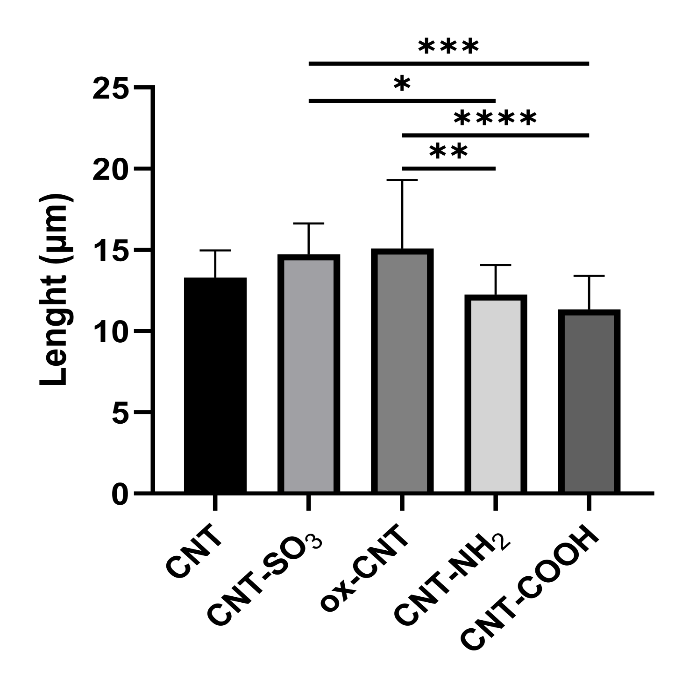

**Figure S1.** Length of nucleus (µm) for each fCNT type (n=20).

|  | Amplitude ±SD | Period ±SD (s) |
| --- | --- | --- |
| Control | 1.228 ±0.150 | 2.154 ± 0.167 |
| pCNT | 1.493 ±0.352 | 2.310 ±1.000 |
| CNT-SO_3_ | 1.154±0.086 | 1.752 ±0,782 |
| ox-CNT | 1.287±0.209 | 1.515 ±0,456 |
| CNT-NH_2_ | 1.891±0.662 | 5.893 ±3.593 |
| CNT-COOH | 1.243±0.226 | 1.723 ±0.705 |

**Table S1.** Amplitude and period average values recorded for all fCNT types.
